# scFormer: A Universal Representation Learning Approach for Single-Cell Data Using Transformers

**DOI:** 10.1101/2022.11.20.517285

**Authors:** Haotian Cui, Chloe Wang, Hassaan Maan, Nan Duan, Bo Wang

**Affiliations:** University of Toronto; Vector Institute; University Health Network; Microsoft Research

## Abstract

Single-cell sequencing has emerged as a promising technique to decode cellular heterogeneity and analyze gene functions. With the high throughput of modern techniques and resulting large-scale sequencing data, deep learning has been used extensively to learn representations of individual cells for downstream tasks. However, most existing methods rely on fully connected networks and are unable to model complex relationships between both cell and gene representations. We hereby propose scFormer, a novel transformer-based deep learning framework to jointly optimize cell and gene embeddings for single-cell biology in an unsupervised manner. By drawing parallels between natural language processing and genomics, scFormer applies self-attention to learn salient gene and cell embeddings through masked gene modelling. scFormer provides a unified framework to readily address a variety of downstream tasks such as data integration, analysis of gene function, and perturbation response prediction. Extensive experiments using scFormer show state-of-the-art performance on seven datasets across the relevant tasks. The scFormer model implementation is available at https://github.com/bowang-lab/scFormer.

## 1 Introduction

Single-cell RNA sequencing (scRNA-seq) is a revolutionary technology that captures gene expression at the resolution of individual cells (Shapiro et al., 2013). Currently, large scRNA-seq atlases already contain tens of millions of cells, and the size of the available data continues to grow exponentially (Regev et al., 2017; Han et al., 2018). This opens up ample opportunities for machine learning algorithms to leverage these large-scale datasets for data-driven discoveries in the field of genomics and medicine.

Recently, deep learning has been employed in the field of single-cell biology to integrate cell embeddings across datasets (Lopez et al., 2018; Gayoso et al., 2021; Lotfollahi et al., 2022), infer cell types (Zhang et al., 2019), analyze gene regulatory networks (Seninge et al., 2021), and predict genetic perturbation responses (Lotfollahi et al., 2019; 2021; Yu & Welch, 2022). The current mainstream model utilized for learning representations of single-cell data are variational autoencoders (VAEs). More specifically, VAEs with multi-layer perceptron (MLP) encoders and decoders are widely used in most of the aforementioned approaches. However, the projection of latent cell embeddings to reconstruct gene expression using MLP layers makes it difficult to model gene-gene interactions, due to the lack of a one-to-one mapping from the compressed latent embedding to the full space of all genes. Yet, we argue simultaneously modelling gene-level and cell-level information is a potential improvement for two reasons: (1) lower-level gene expression and higher-level cell identity are highly correlated, and the embeddings can share information during model training; (2) single-cell analysis focuses on both cell and gene related tasks. Common Cell-level tasks include cell annotation and clustering, and gene-level tasks include functional pathway enrichment and gene network analysis. Therefore, the capability to provide both cell and gene representations is a desirable property of models that lead to applicability to multiple downstream tasks.

To simultaneously provide gene and cell representations, we propose scFormer, a transformer-based model that utilizes self-attention on gene expression and provides jointly optimized cell and gene embeddings. Recently, unsupervised learning of large datasets using self-attention transformers (Vaswani et al., 2017) has shown major success in several machine learning fields, including language (Devlin et al., 2018; Brown et al., 2020), computer vision (He et al., 2022), and learning protein representations (Jumper et al., 2021; Rao et al., 2021; Baek et al., 2021). Similarly, we argue that the self-attention model can readily be applied to sequencing data and learn the context-specific correlated expression patterns in an unbiased way. In this work, (1) We introduce several techniques including masked gene modeling (MGM), and masked value for cell (MVC), to facilitate self-attention optimization in the single-cell domain. (2) To our best knowledge, scFormer is the first work using transformer to jointly learn cell and gene embeddings in a completely unsupervised fashion and attain representations applicable to multiple downstream tasks. We envision scFormer to be a new backbone model for single cell modeling due to these advantages, and we show the techniques promising performance across several downstream tasks on seven datasets.

## 2 Related Work

### Learning cell and gene representation for scRNA-seq

Cell representation learning has been one of the major research areas in the field of single-cell data modelling. Cell embeddings are the foundation for various downstream tasks, such as cell type annotation, visualization, and data integration. Early approaches like Seurat (Satija et al., 2015) and its variants (Stuart et al., 2019) adopt nearest-neighbor-based alignment to correct technical effects (Eisenstein, 2020) and learn linearly transformed cell embeddings. Other approaches like LIGER (Liu et al., 2020) and OCAT (Wang et al., 2022) use matrix factorization to find robust cell embeddings. Aside from these techniques, Deep Learning methods, particularly VAE-based generative models (Kingma & Welling, 2013), have been widely used in recent studies. scVI (Lopez et al., 2018) learns latent cell representations by variational inference and reconstructing original gene expressions. TotalVI (Gayoso et al., 2021) and scGen (Lotfollahi et al., 2019) utilized similar models and extended the applications to multi-omics and perturbation prediction. On the other hand, gene representation learning benefits many downstream tasks, including gene regulatory network and functional pathway analysis. As an example, GeneVector (Ceglia et al., 2022) detects gene-gene functional relations by factorizing the co-expression and mutual information matrix of the sequencing readout. Despite the importance of the two branches of research for cell and gene embeddings, few approaches have worked on jointly learning both. DeepMAPS (Ma et al., 2021) utilizes graph neural networks to encode cell and gene nodes for related tasks. scFormer stands out as an approach to effectively learn both embeddings of cells and genes jointly in a shared architecture.

### Transformers for modelling scRNA-seq data

Transformer models with self-attention (Vaswani et al., 2017) have achieved great success in natural language processing (NLP) (Devlin et al., 2018), and recently in computer vision and protein biology as well. Despite these results, there have been few attempts to adopt the transformer architecture into single-cell biology and applications thereof. Shen et al. (2022) use the transformer decoder setup to learn the gene name sequence of highly expressed genes, without considering the actual sequenced expression abundance. This leads to loss of major biological signal, as the expression values are informative of cell state and gene-gene relationships. A very recent work, scBERT (Wang et al., 2021), that uses a BERT-like architecture (Devlin et al., 2018) only applied learned embeddings for the supervised task of cell annotation. To our best knowledge, scFormer is one of the first methods to provide a transformer backbone for multiple single-cell analysis tasks in an unsupervised fashion.

## 3 Methods

Single-cell sequencing captures genetic sequence information from individual cells, in contrast to bulk approaches that average information across many cells. In particular, the widely used scRNA-seq measures the individual abundance of RNA molecules in each cell, providing a profile of the cellular identity, stage, and functionality. The scRNA-seq transcriptomic data is quantified into a cell-gene matrix, 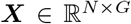, where each entry 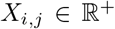 is the read for the transcribed RNA abundance of gene *j* ∈ {0,1,…, *G*} in the cell *i* ∈ {0,1,…, *N*}. We refer to this data as the raw matrix in later sections. Different from natural language texts, the sequenced data are continuous and therefore create challenges for tokenization and modelling. Next, we introduce both data processing techniques and learning objectives to facilitate the learning using transformer.

### 3.1 Input embeddings

The input to scFormer consists of three components: (1) gene tokens, (2) gene expression values, and (3) optional external tokens. The gene tokens and expression values are pre-processed from the raw matrix X with a slightly different procedure for each modeling task (see section 4).

#### Gene Tokens

We naturally use gene names as gene tokens. Each gene *g_j_* has a unique integer id *id*(*g_j_*) out of the full vocabulary of tokens. Notably, for a dataset of multiple studies that contain different sets of genes due to technological/processing differences, these tokens can be readily collected into a shared vocabulary of the union set of genes across studies. This leads to a unique flexibility of scFormer for modeling data with distinct gene sets when integrating multiple studies. We also include special tokens in the vocabulary, including < *cls* > for integrating across genes into a cell representation and < *pad* > for padding the input length in a mini-batch. Conceptually, we consider the gene tokens work similar to the word/token embeddings in natural language modeling (NLM). In summary, the input gene tokens for each cell *i* are a fixed length vector 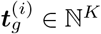,

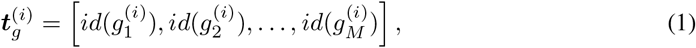

where *M* is the preset input length, which is usually set as the number of highly variable genes used.

#### Expression Values

The input of gene expression values are converted from the raw counts *X_i,j_*. A key challenge of modeling gene expression is that the absolute magnitudes vary among sequencing protocols (Sarkar & Stephens, 2021). Because of the difference in sequencing depth and in probability of capturing lowly expressed genes, data from different sequencing batches (the terminology for experiment trials) have quite different scales even after common preprocessing measures of normalizing to a fixed sum and *log*1*p* transformation. In other words, the same absolute value conveys different *“semantic”* meaning across sequencing batches. To resolve this, we introduce ***value binning*** and convert all expression counts into relative values. For all non-zero expression counts of each cell, we count the raw absolute values and make *B* number of consecutive intervals [*b_k_*,*b*_*k*+1_],*k* ∈ {1, 2,…, *B*}, where each interval range includes an equal 1/*B* portion of all expressed genes. Note that the computation is done cell-wise and the interval edges *b_k_* vary among cells. The converted expression value 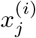 for each cell *i* is as follows,

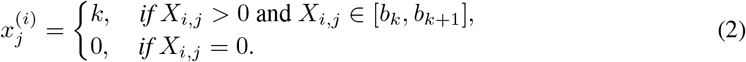

With this binning, 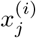 has consistent semantic meaning across sequencing batches. For example, the value 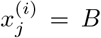 always implies that gene *j* is one of the highest expressed genes in the cell. Conventional pre-processing steps Luecken & Theis (2019) are conducted before the value binning step. Here we omit these other steps and use the raw data notation *X_i,j_* in the above equation. The final input value vector for cell *i* is

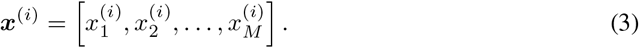

#### External Tokens

The external tokens can contain any meta information corresponding to individual genes. For example, pathway tokens can represent the functional pathways a gene belongs to, and perturbation tokens can indicate if a gene is altered in perturbation experiments. We describe all external tokens as an input vector with the same dimension as the input genes,

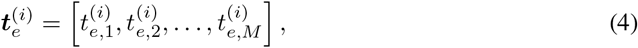

where 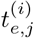 are integer indices representing external categories.

#### Embedding layers

We use standard embedding layers^1^ emb_g_ for the gene tokens and emb_e_ for the external tokens to map each token into a embedding vector of fixed-length *D*. Although the standard embedding layer is also applicable to the expression values since they are binned into fixed set of *B* + 1 integers, we by default use fully connected layers, emb_x_. This has the benefit of easily modeling the consecutive nature of the value magnitudes. The final embedding 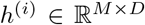 of cell *i* is defined as

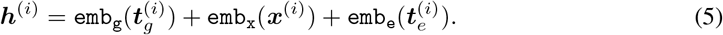

### 3.2 Encoder and gene expression modeling

We use the transformer encoder (Vaswani et al., 2017; Devlin et al., 2018) to encode the total input embedding *h*^(*i*)^ in equation 5. The self-attention mechanism in transformer blocks operates over the *M* embedding vectors in the input sequence, and particularly suits the goal of learning the interaction between genes across different cell types. The output of stacked transformer blocks is

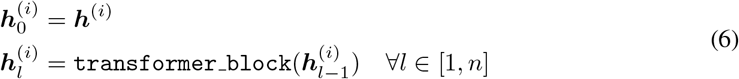

We use the output, 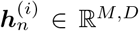, for both gene-level and cell-level tasks. Gene-level task heads (see section 3.4) can be directly applied on these learned embeddings for specific tasks including perturbed expression prediction and masked gene modelling. For cell-level tasks, we first integrate 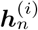 into a cell embedding vector (section 3.3).

The input number *M* of genes can go up to tens of thousands. This greatly exceeds the input length of common transformers used in NLM. Efficient self-attention techniques can be used (Katharopoulos et al., 2020; Wang et al., 2020; Dao et al., 2022). Also, since the order of the genes is not sequential in scRNA-seq data, and the transformer computation is agnostic to the order, we can dynamically sample subsets of the input.

### 3.3 Cell representation

We view each cell as a “sentence” of genes, and a cell representation 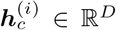 vector can be generated by integrating the learned gene-level representation 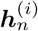. Common pooling operations such element-wise mean-pooling or weighted-pooling can be readily used. Here, we choose to use a special token < *cls* > for the cell representation and let the model learn the pooling operation within transformer blocks: the < *cls* > token is appended to the beginning of other input tokens (figure 1), and the final embedding at this position will be extracted as the cell representation, which is usually the first row of 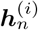 i.e. 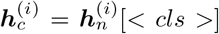, where [< *cls* >] denotes retrieving the row at the index of < *cls* > token in the input.

**Figure 1:**
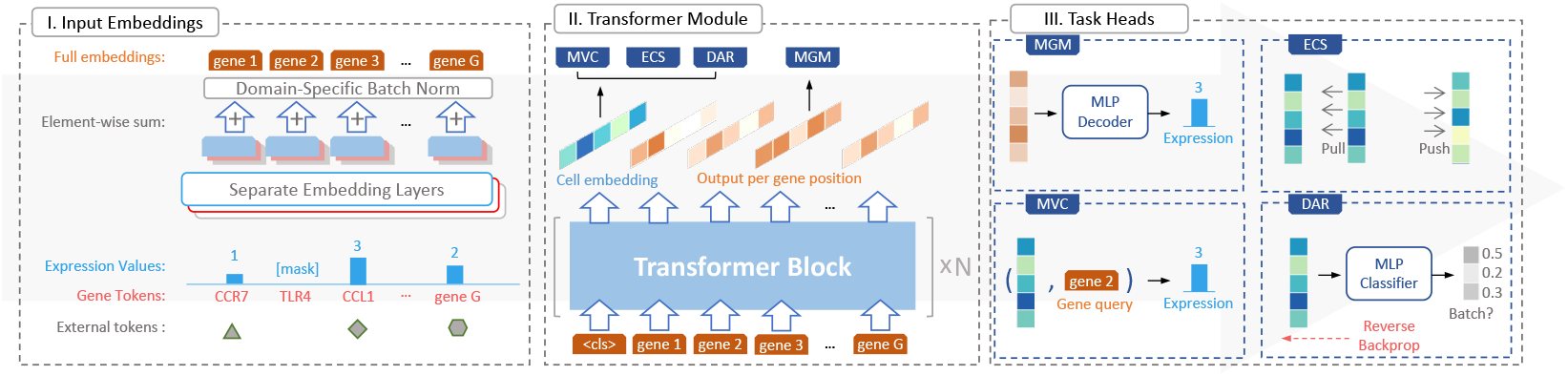
Model schematic. From left to right: I. input embeddings that integrate three types of tokens. II. the stacked transformer encoder blocks. III. Four task heads that are applied on cell embeddings and gene-level transformer output.

### 3.4 Task heads

We use the term task head for the task-specific module and the loss function paired with it. scFormer benefits from various task heads to facilitate the learning of biologically meaningful cell and gene representations, in addition to task heads for regularization purposes such as batch correction.

#### Masked Gene Modelling (MGM)

scFormer employs masked gene modelling to promote the learning of cross-gene relations, inspired by the masked-language modelling in NLM. In each cell, a proportion of genes and their expression values *x*^(*i*)^ are randomly masked, and the scFormer model is optimized to correctly predict the gene expression values at the masked positions. This task head helps the model effectively encode co-expression within sets of genes. Formally, we feed the transformer output into a fully connected MLP to estimate the expression value for *M* genes, and use the cross entropy loss (ce) only at the masked positions, ℳ_*mask*_, to optimize this objective:

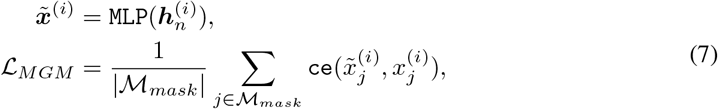

where 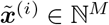 is the row of expression estimates.

MGM is a general self-supervised task head that works to predict gene expression for masked genes. In some downstream tasks, such as the perturbation prediction task, the model will predict known target gene expression values rather than the original ones. In such a supervised scenario, no masking is needed. We keep the MLP estimator and cross entropy loss in equation 7, use target gene expression as 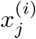 in the equation, and simply change the predicted expression values to apply to all valid target positions instead of the masked positions.

#### Masked Value for Cell Modelling (MVC)

This task head works in a similar fashion as the MLM, although it instead uses and promotes the cell representation 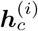. For the expression of each gene *j* in an input cell *i*, we make a query vector *q_j_* and use the parameterized inner product of *q_j_* and cell representation 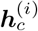 as the predicted expression value.

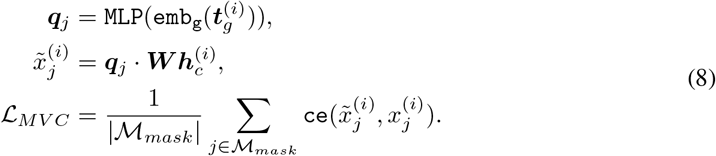

MVC shares the the gene token embedding, 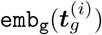 in equation 5. In practice, we found applying MGM and MVC altogether achieves significantly better performance than applying either individually (section 4.2.1). This is consistent with our argument that the joint modeling of cell and gene representations contributes to learning a more biologically meaningful embedding of both.

#### Elastic Cell Similarity (ECS)

This task head enhances the disentangling of cell representations. It uses a contrastive learning loss introduced by Liu et al. (2019).

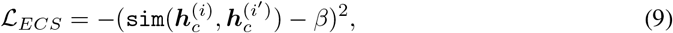

where `sim` is the cosine similarity function, i, i′ denote two cells in the same training mini-batch, and *β* is a predefined threshold. ECS works pair-wise on all cells in a mini-batch. Intuitively, it increases the similarity of the pairs that already have similarity above *β*, and conversely pushes away dissimilar pairs.

#### Domain Adaptation by Reverse Back-propagation (DAR)

Technical batch effects introduce non-biological differences between data samples and can greatly impact the representation learning procedure (Eisenstein, 2020; Tran et al., 2020). To address this issue, we use a separate MLP classifier to predict the sequencing batch of each input cell, and reverse the gradients when back-propagating through the classifier. This strategy has been shown as a robust domain adaptation method by Ganin & Lempitsky (2015). We also use a domain-specific batch normalization (DSBN) (Chang et al., 2019) on the input embedding (equation 5) as a soft strategy to further enhance batch correction.

## 4 Experiments and results

### 4.1 Representation Learning For single dataset of scRNA-seq

Cell clustering and visualization is an essential task that often serves as the first step for cell type and cell state identification. Therefore, we first evaluate the clustering on datasets with annotated cell types to measure how well the cell representations learned by scFormer preserve biological information as such.

#### Datasets

We tested 3 datasets re-procecessed by Gayoso et al. (2022): Cortex (3,005 cells and 19,972 genes), PBMC 8K (7,982 cells and 3,346 genes), and Spleen 17K (17,001 cells and 13,553 genes).

#### Experiment Setup

We performed the following preprocessing steps using the SCANPY python library (Wolf et al., 2018): (1) normalize each cell by total counts over all genes, (2) logarithmize the data matrix with log1p, and (3) select highly variable genes.

We evaluated cell embeddings on biological conservation metrics proposed in Luecken et al. (2022). Biological conservation evaluation metrics included *NMI* (normalised mutual information), *ARI* (adjusted rand index), and *ASW* (average silhouette width), to measure the consistency between derived cell type clusters and ground truth labels. For ease of comparison, we also reported *AvgBIO* as the average of *NMI, ARI* and *ASW_cell_*. See Appendix A.1 for details on metric calculations.

We benchmarked scFormer against Seurat (Satija et al., 2015), scVI (Lopez et al., 2018) and a highly variable gene (HVG) baseline on all datasets. For all methods benchmarked, we used the same set of highly variable genes. The output metrics are calculated using the implementation in `scib.metrics` by Luecken etal. (2022).

#### Results

The benchmark in single datasets demonstrates that scFormer achieves the state-of-the-art results in the cell embedding extraction task across all metrics tested. Notably, scFormer’s *AvgBIO* score on the Cortex dataset exceeds other methods by a 8–17% margin, and on the PBMC 8K dataset by a 5 – 19% margin. This showcases scFormer’s superior performance in amplifying biological and cell type signals through effective feature learning. See Appendix A.2 for the comparisons of UMAP visualizations on the Cortex dataset.

### 4.2 Integration of multiple scRNA-seq data with batch correction

Cell representation learning faces the challenge of batch effects when multiple datasets or sequencing batches are given as input. True biological variance may be confounded with technical difference between input batches. Without batch correction, two cells from the same batch of different cell types maybe be clustered together rather than two of same cell type from different batches, leading to errors in cell type annotations. Therefore, we assess scFormer’s ability to correct batch effects while preserving biological variance of the integrated datasets.

#### Datasets

For data integration task, we tested on 2 datasets re-processed by (Gayoso et al., 2022) and (Luecken et al., 2022): Immune human (33,506 cells and 12,303 genes from 10 donors), and Pancreas (16,382 cells and 19,093 genes from 9 batches).

#### Experiment Setup

In integration datasets, we performed the same preprocessing steps as described in Section 4.1. Additionally, we filtered out genes with low read counts as quality control suggested by Luecken et al. (2022).

We reported the same biological conservation metrics and *AvgBIO* for cell type clustering as described in Section 4.1. Additionally, we reported batch correction metrics proposed in Luecken et al. (2022) to assess batch mixing. Batch correction performance is measured by *ASWbatch*, the inverse of average silhouette width for batch clustering, and *GraphConn* for graph connectivity. For ease of comparison, we reported *AvgBATCH* as the average of *ASWbatch* and *GraphConn* for batch mixing. We also reported an *Overall* score as a weighted sum of *AvgBIO* and *AvgBATCH*, consistent with Luecken et al. (2022). See Appendix A.1 for details on metric calculations.

We benchmarked scFormer against other unsupervised methods, including Seurat (Satija et al., 2015), Harmony (Korsunsky et al., 2019), and scVI (Lopez et al., 2018). Harmony and scVI has been shown with best performances in recent benchmarking of integration methods Luecken et al. (2022). For all methods benchmarked, we used the same set of highly variable genes across all methods.

#### Results

On both datasets, scFormer achieves the best biological conservation score (*AvgBIO*) and the best overall score. Shown in figure 2a, we find the learned cell representation can be well clustered in concordance with the cell type labels. scFormer also provides comparable batch correction results (figure 2b), although the performance is not fully shown in the *AvgBATCH* score (see section 4.2.1). Note that the biological conservation is a more important evaluation for this task, since it ensures that the clusters are reliable for downstream analysis such as cell type annotation.

**Figure 2:**
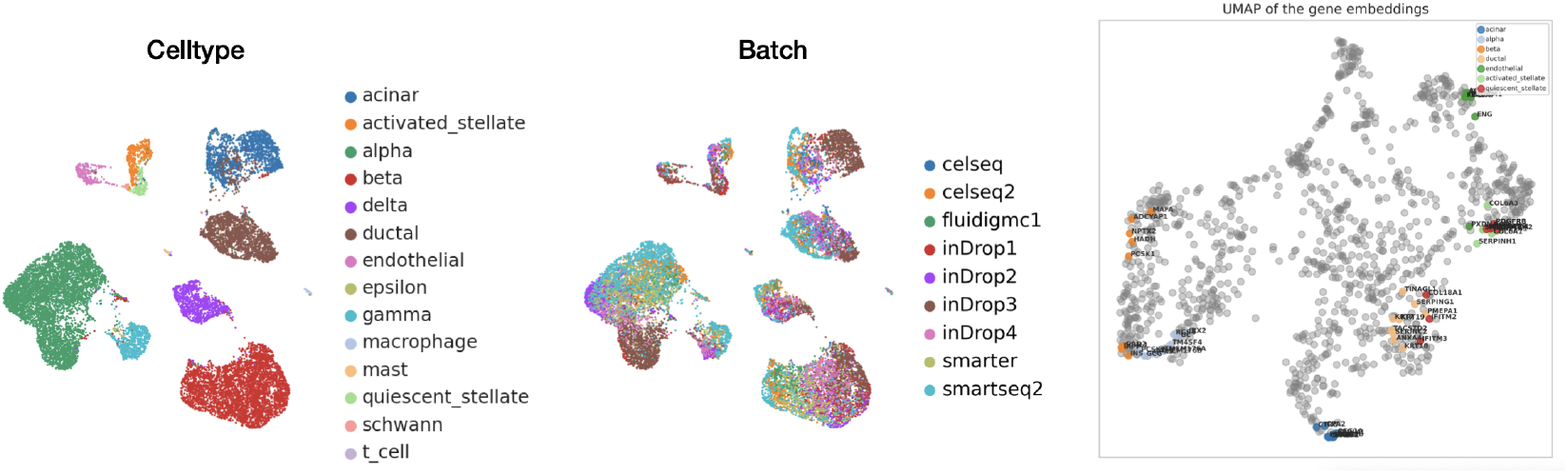
(*a*)UAMP plot of learned gene embeddings with colors of *cell types*. (*b*)UAMP plot of learned gene embeddings with colors of *sequencing batches*. (*c*)UMAP of gene embedding. Colored genes are top 9 markers of corresponding cell types. All other genes in the input space are shown in grey.

Using both MVC and MGM task heads, scFormer learns gene embeddings simultaneously with cell embeddings. We present the 2-dimensional UMAP (McInnes et al., 2018) plot of the learned gene embeddings of 1200 input genes. As shown in Figure 2, the gene markers of major cell types are well clustered together, even though scFormer is trained in a pure unsupervised manner without cell type labels. These results demonstrate that scFormer is capable of simultaneously learning distinguishable cell and gene representation.

#### 4.2.1 Ablation studies

We validated the effectiveness of each task head through ablation experiments on the Pancreas dataset, as detailed in Table 3. In the ablation study, we tested 9 options including: (1) full scFormer with all task heads on and default MGM mask ratio, (2) 2 additional full task-headed scFormer with lower or higher MGM mask ratios, and (3) 6 ablation settings with removal of one or two task heads. Each option is repeated 5 times for random seed values 0-4, and the best cell clustering and batch mixing metrics are reported.

**Table 1:** Cell embedding results (Single dataset)

| Model | Cortex |  |  |  | PBMC 8K |  |  |  | Spleen 17K |  |  |  |
| --- | --- | --- | --- | --- | --- | --- | --- | --- | --- | --- | --- | --- |
|  | AvgBIO | ARI | NMI | ASW<sub>cell</sub> | AvgBIO | ARI | NMI | ASW<sub>cell</sub> | AvgBIO | ARI | NMI | ASW<sub>cell</sub> |
| HVG | 0.605 | 0.657 | 0.643 | 0.517 | 0.626 | 0.624 | 0.733 | 0.522 | 0.600 | 0.611 | 0.685 | 0.505 |
| Seurat | 0.597 | 0.618 | 0.622 | 0.551 | 0.762 | **0.864** | 0.821 | 0.601 | 0.641 | **0.650** | 0.702 | 0.572 |
| scVI | 0.688 | 0.743 | 0.716 | 0.606 | 0.696 | 0.699 | 0.797 | 0.592 | 0.619 | 0.615 | 0.692 | 0.551 |
| scFormer | **0.763** | **0.805** | **0.738** | **0.745** | **0.819** | 0.786 | **0.861** | **0.810** | **0.645** | 0.640 | **0.706** | **0.589** |

**Table 2:** Cell embedding results (Integration)

| Model | Immune Human |  |  | Pancreas |  |  |
| --- | --- | --- | --- | --- | --- | --- |
|  | AvgBIO | AvgBATCH | Overall | AvgBIO | AvgBATCH | Overall |
| Seurat | 0.565 | 0.882 | 0.691 | 0.647 | 0.910 | 0.752 |
| Harmony | 0.743 | 0.914 | 0.811 | 0.836 | 0.916 | 0.868 |
| scVI | 0.725 | **0.921** | 0.803 | 0.829 | **0.917** | 0.864 |
| scFormer | **0.765** | 0.903 | **0.820** | **0.882** | 0.900 | **0.889** |

**Table 3:**
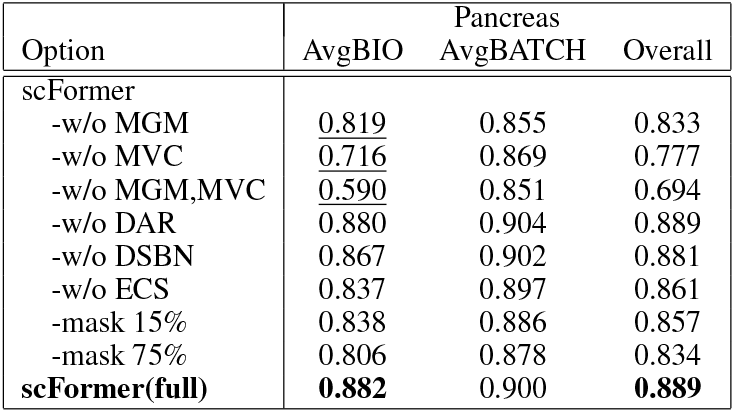
Ablation Results.

Notably, MGM and MVC task heads are critical in learning biological information, as the model observes a 6 – 16% drop in AvgBIO when one is removed, and 29% drop when both are removed. See Figure 3 for much improved cluster separation by cell type and mixing by batch: for example, intertwined cell types alpha and gamma become distinctly separated, and the three batch clusters for cell type beta have merged into one. DAR, ECS, and DSBN task heads all contributed to the smoothing effects on batch mixing, despite comparable scores in the *ASW_batch_* metric. Furthermore, the appropriate mask ratio is essential for effective learning, as demonstrated in the performance deterioration when mask ratio is too high or two low.

**Figure 3:**
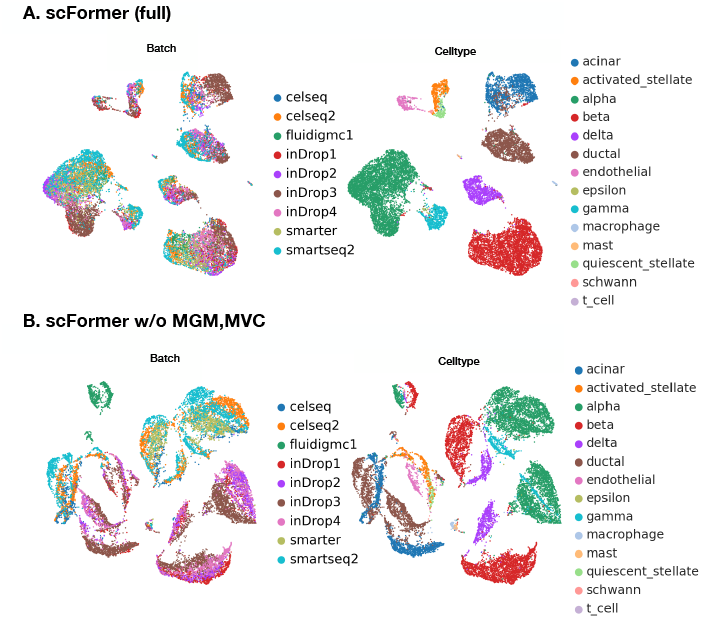
UMAP Visualizations.

### 4.3 Perturbation Prediction

Recent combination of scRNA-seq and gene editing techniques enables high-throughput experiments revealing the cellular response to multiple genetic perturbations. This has become a promising tool for the discovery of novel gene interactions and regenerative medicine. However, the combinatorial space of possible gene being perturbed quickly exceeds the scope of feasible experiments and thus limits the application. Therefore, machine learning methods can be applied to learn from cellular response of known experiments and extrapolate to unknown ones. scFormer is particularly suitable for this task because the self-attention over gene dimension may well encoded the interaction between perturbed genes and downstream expression responses of other genes. We test scFormer in this setting of predicting gene expressions after perturbation and shows its performance.

#### Datasets

For perturbation task, we benchmarked on 2 perturbation datasets pre-processed by Roohani et al. (2022): (1) Pertub-seq dataset by Adamson et al. (2016) containing 87 1-gene perturbations, with around 100 cells per perturbation and at least 7,000 unperturbed cells, and (2) Perturb-Seq dataset by Norman et al. (2019) containing 131 2-gene perturbations and 105 1-gene perturbations, with 300-700 cells treated with each perturbation.

#### Experiment Setup

We followed the same preprocessing steps by Roohani et al. (2022) in their benchmark: (1) normalize by total counts over all genes, (2) log transform data, (3) select 5000 highly variable genes, and (4) include any pertubed genes not accounted for. In our experiments, for 1-gene perturbation prediction in both datasets (Adamson et al., 2016; Norman et al., 2019), the perturbations are split to ensure that test perturbations are not seen in training, i.e., no cells in training set has undergone any of the test perturbations. For 2-gene perturbation prediction in the Norman et al. (2019) dataset, the train-test split consists of three scenarios with increasing difficulty: (1) 0/2 unseen genes, (2) 1/2 unseen genes, and (3) 2/2 unseen genes in the training set.

We evaluate perturbation prediction accuracy based on Pearson correlation (*corr*) between predicted gene expressions post-perturbation and ground-truth expression values. Another variant of the Pearson metric is calculated on the amount of change in expression post-perturbation compared to control instead of raw expression values, denoted as *corr*(Δ). We also report these Pearson metrics on different gene sets, including (1) all genes (*ALL*), and (2) top 20 differentially expressed genes (*DE*). We thus report 4 evaluation metrics as detailed below, namely *corr* and *corr*(Δ) each for gene sets (*ALL*) and (*DE*). See Appendix A.1 for details on metric calculation.

#### Results

We compare the performance against the recent GEARS method (Roohani et al., 2022) and the multi layer perceptron baseline. scFormer shows the highest correlation to the ground-truth perturbed expressions on almost all metrics. Since around 50% of gene expression counts before and after perturbation are zero due to either low capture rate or low expression, we would argue that the evaluation on differentially expressed genes, i.e., the DE columns in Table 4, are more convincing. Particularly, significant improvements by scFormer are shown for the correlation of the change (Δ) of the top differentially expressed genes, which is arguably the most important metric.

**Table 4:** Perturbation generation results

| Model | Norman et al. (2019) |  |  |  | Adamson et al. (2016) |  |  |  |
| --- | --- | --- | --- | --- | --- | --- | --- | --- |
|  | DE |  | ALL |  | DE |  | ALL |  |
| | corr | corr($\Delta$) | corr | corr($\Delta$) | corr | corr($\Delta$) | corr | corr($\Delta$) |
| MLP | 0.909 | 0.428 | 0.987 | 0.408 | 0.948 | 0.729 | 0.991 | **0.656** |
| GEARS | 0.917 | 0.508 | 0.986 | 0.387 | 0.961 | 0.726 | 0.991 | 0.652 |
| scFormer | **0.921** | **0.582** | **0.988** | **0.459** | **0.964** | **0.740** | **0.991** | 0.632 |

## 5 Implementation details

All models are set to have 4 stacked transformer blocks. Each block has an embedding size of 128, 4 attention heads and the fully connected layer has hidden size of 128. The training mini-batch is set to 16. We use the Adam optimizer with a starting learning rate 0.001, and decay to 90% after each epoch. We set the mask ratio of MGM and MVC to 0.4, *beta* in ECS to 0.6, and a weighting of 10 on ECS loss when combined with others. For the embedding learning tasks (sections 4.1 and 4.2), each dataset is split into train and evaluation sets at 9:1 ratio. We trained the model for fixed 30 epochs and evaluated the MGM loss value on the validation set after each epoch. We report the model with the best validation score. For the perturbation task (section 4.3), we noticed the model can converge usually within 3 epochs and we similarly report the best validated model.

## 6 Conclusion

We hereby propose scFormer, a novel transformer-based deep learning framework to jointly optimize cell and gene embeddings for single-cell biology in pure unsupervised manner. scFormer provides a unified framework to address a variety of downstream tasks including data integration, gene function analysis, and perturbation response prediction. Empirical results show that the self-attention on gene expressions and the introduced MGM and MVC objectives significantly boost the performance for cell-level and gene-level tasks. For future directions, we envision the proposed techniques can be applied to other modalities such as single-cell atac-seq and spatial transcriptomics.

## A Appendix

### A.1 Evaluation Metric Calculations

#### A.1.1 Embedding Extraction

We followed the evaluation metric calculations specified by Luecken et al. (2022) in their benchmark paper as detailed below.

##### Normalized Mutual Information

We calculate the normalized mutual information (NMI) score to measure the overlap between ground truth cell type labels and Louvain cluster labels obtained from integrated cell embeddings. Louvain clustering was performed at a resolution range of 0.1 to 2 in steps of 0.1 to identify the highest NMI to be reported. The cell type NMI score, denoted as ***NMI_cell_***, ranges from 0 to 1, with higher score indicating better cell type match.

##### Adjusted Rand Index

We calculate the adjusted rand index (ARI) to measure both overlap and disagreements between ground truth cell type labels and MNI-optimized Louvain clusters. The rand index is further adjusted for randomly correct labels. The cell type ARI score, denoted as ***ARI_cell_***, ranges from 0 to 1, with 0 corresponding to random labelling and 1 for perfect match.

##### Average Silhouette Width

The silhouette width measures the relationship between the within-cluster distances of a cell and the between-cluster distances of that cell to the closest cluster. The average silhouette width (ASW) score is calculated by averaging the silhouette widths of all cells. The ASW score ranges from −1 and 1, where an ASW score of 1 suggests well-separated clusters while −1 to 0 implies overlapping clusters and misclassification.

For cell type clustering evaluation, we calculate the ASW score with respect to cell type labels, denoted as ***ASW_cell_***:

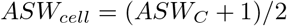

where *C* denotes cell types.

For batch mixing evaluation, we calculate the ASW score with respect to batch labels and scale it by subtracting 1, denoted as ***ASW_batch_***:

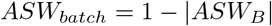

Both ***ASW_cell_*** and ***ASW_batch_*** range from 0 to 1, with higher score indicating better cell type clustering or batch mixing performance.

##### Graph Connectivity

The graph connectivity metric computes the average proportion of cells that are connected through a kNN graph within its own cell type. For each cell identity *c* in *C*, we calculate the size of the largest connected component with kNN among cells of identity *c* only over the total number of cells of identity *c*. The average across all cell types is reported as the ***GraphConn*** score:

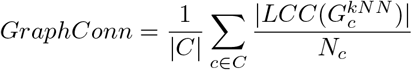

where *LCC* denotes the largest connected component and *N* denotes total number of cells for each cell type.

##### Aggregated Metrics

The aggregated metric ***AvgBIO*** computes the mean of the three metrics for biological conservation:

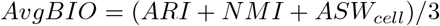

The aggregated metric ***AvgBATCH*** computes the mean of the two metrics for batch mixing:

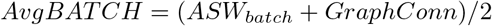

Following the convention in (Luecken et al., 2022), an ***Overall*** metric for integration tasks is computed as the weighted average of ***AvgBIO*** and ***AvgBATCH***:

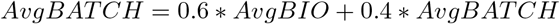

#### A.1.2 Perturbation Prediction

##### Pearson Correlation

Pearson Correlation (*corr*) is used to measure the correlation between the mean predicted expressions and mean ground truth expressions for the perturbation group. Similarly, *corr*(Δ) computes the correlation on change in the mean expressions post-perturbation compared to control. The Pearson metric is calculated using `scikit-learn`’s implementations.

#### A.2 Cell Embedding Task Results - UMAP Visualizaions

**Table 5:** Integration metrics details (Immune Human)

| Model | Biological Conservation |  |  |  | Batch Mixing |  |  |
| --- | --- | --- | --- | --- | --- | --- | --- |
|  | AvgBIO | ARI | NMI | ASW<sub>cell</sub> | AvgBATCH | ASW<sub>batch</sub> | GraphConn |
| Seurat | 0.565 | 0.445 | 0.695 | 0.556 | 0.882 | 0.858 | 0.907 |
| Harmony | 0.743 | 0.830 | 0.810 | 0.590 | 0.914 | 0.860 | 0.968 |
| scVI | 0.725 | 0.780 | 0.813 | 0.582 | **0.921** | **0.871** | 0.971 |
| scFormer | **0.765** | **0.844** | **0.821** | **0.632** | 0.903 | 0.832 | **0.975** |

**Table 6:** Integration metrics details (Pancreas)

| Model | Biological Conservation |  |  |  | Batch Mixing |  |  |
| --- | --- | --- | --- | --- | --- | --- | --- |
|  | AvgBIO | ARI | NMI | ASW<sub>cell</sub> | AvgBATCH | ASW<sub>batch</sub> | GraphConn |
| Seurat | 0.647 | 0.557 | 0.769 | 0.616 | 0.910 | 0.841 | **0.980** |
| Harmony | 0.836 | 0.94 | 0.91 | 0.66 | 0.916 | **0.880** | 0.952 |
| scVI | 0.829 | 0.949 | 0.914 | 0.625 | **0.917** | 0.863 | 0.972 |
| scFormer | **0.882** | **0.954** | **0.921** | **0.773** | 0.900 | 0.833 | 0.968 |

**Table 7:**
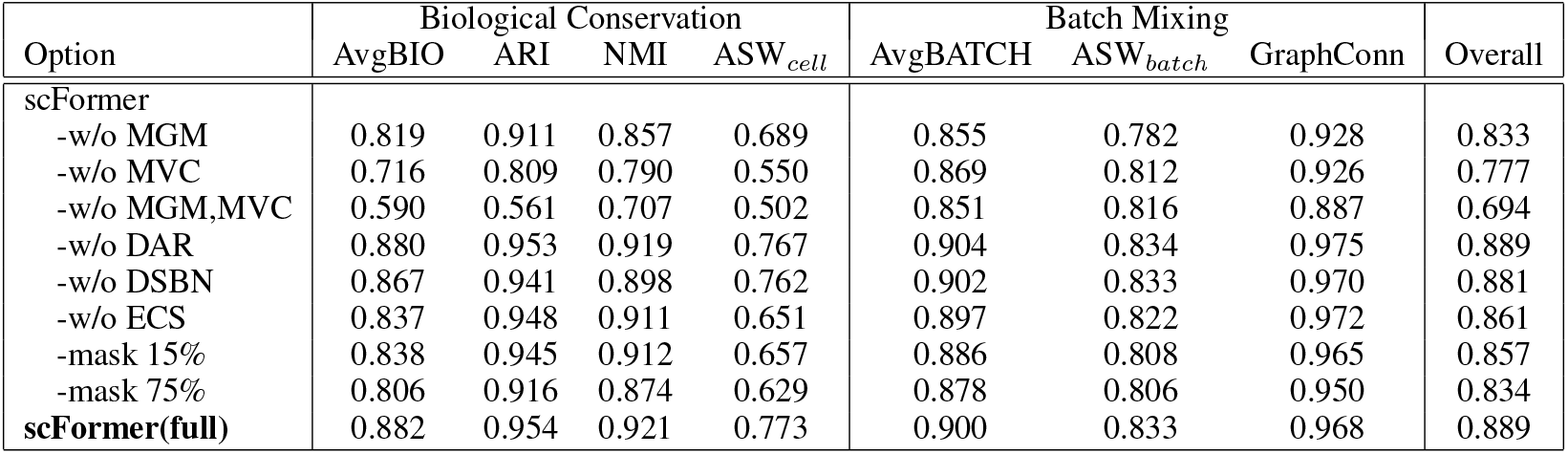
Ablation metric details (Pancreas)

| Option | Biological Conservation |  |  |  | Batch Mixing |  |  | Overall |
| --- | --- | --- | --- | --- | --- | --- | --- | --- |
|  | AvgBIO | ARI | NMI | ASW<sub>cell</sub> | AvgBATCH | ASW<sub>batch</sub> | GraphConn |  |
| scFormer |  |  |  |  |  |  |  |  |
| -w/o MGM | 0.819 | 0.911 | 0.857 | 0.689 | 0.855 | 0.782 | 0.928 | 0.833 |
| -w/o MVC | 0.716 | 0.809 | 0.790 | 0.550 | 0.869 | 0.812 | 0.926 | 0.777 |
| -w/o MGM,MVC | 0.590 | 0.561 | 0.707 | 0.502 | 0.851 | 0.816 | 0.887 | 0.694 |
| -w/o DAR | 0.880 | 0.953 | 0.919 | 0.767 | 0.904 | 0.834 | 0.975 | 0.889 |
| -w/o DSBN | 0.867 | 0.941 | 0.898 | 0.762 | 0.902 | 0.833 | 0.970 | 0.881 |
| -w/o ECS | 0.837 | 0.948 | 0.911 | 0.651 | 0.897 | 0.822 | 0.972 | 0.861 |
| -mask 15% | 0.838 | 0.945 | 0.912 | 0.657 | 0.886 | 0.808 | 0.965 | 0.857 |
| -mask 75% | 0.806 | 0.916 | 0.874 | 0.629 | 0.878 | 0.806 | 0.950 | 0.834 |
| **scFormer(full)** | 0.882 | 0.954 | 0.921 | 0.773 | 0.900 | 0.833 | 0.968 | 0.889 |

## Footnotes

1 look-up table embedding layer, https://pytorch.org/docs/stable/generated/torch.nn.Embedding.html

